# Different outcomes of endurance and resistance exercise in skeletal muscles of Oculopharyngeal muscular dystrophy

**DOI:** 10.1101/2024.01.12.575335

**Authors:** Alexis Boulinguiez, Jamila Dhiab, Barbara Crisol, Laura Muraine, Ludovic Gaut, Corentin Rouxel, Justine Flaire, Hadidja-Rose Mouigni, Mégane Lemaitre, Benoit Giroux, Lucie Audoux, Benjamin SaintPierre, Arnaud Ferry, Vincent Mouly, Gillian Butler-Browne, Elisa Negroni, Alberto Malerba, Capucine Trollet

**Affiliations:** Department of Biological Sciences, School of Life Sciences and the Environment, Royal Holloway University of London, Egham, Surrey TW20 0EX, UK; Myology Center for Research, U974, Sorbonne Université, INSERM, AIM, GH Pitié Salpêtrière Bât Babinski, 47 bd de l’Hôpital, 75013, Paris, France; Sorbonne Université, INSERM, UMS28 – Phénotypage du petit animal, 105 Blvd de l’Hôpital Paris 75013; Université Paris Cité, CNRS, INSERM, Institut Cochin, F-75014 Paris, France

**Keywords:** Skeletal muscle, OPMD, PAPBN1, Exercise, Atrophy, Fibrosis

## Abstract

**Background:** Exercise is widely considered to have beneficial impact on skeletal muscle aging. In addition, there are also several studies demonstrating a positive effect of exercise on muscular dystrophies. Oculopharyngeal muscular dystrophy (OPMD) is a late-onset autosomal dominant inherited neuromuscular disorder caused by mutations in the *PAPBN1* gene. These mutations consist in short (1-8) and meiotically stable GCN trinucleotide repeat expansions in its coding region responsible for the formation of PAPBN1 intranuclear aggregates. This study aims to characterize the effects of two types of chronic exercise, resistance and endurance, on the OPMD skeletal muscle phenotype using a relevant murine model of OPMD.

**Methods:** In this study, we tested two protocols of exercise. In the first, based on endurance exercise, FvB (wild-type) and A17 (OPMD) mice underwent a 6-week-long motorized treadmill protocol consisting in 3 sessions per week of running 20cm/s for 20 minutes. In the second protocol, based on resistance exercise generated by chronic mechanical overload (OVL), surgical removal of gastrocnemius and soleus muscles was performed, inducing hypertrophy of the plantaris muscle. In both types of exercise, muscles of A17 and FvB mice were compared to those of respective sedentary mice. For all the groups, force measurement, muscle histology and molecular analyses were conducted.

**Results:** Following the endurance exercise protocol, we did not observe any major changes in the muscle physiological parameters, but an increase in the number of PABPN1 intranuclear aggregates and enhanced collagen deposition in the exercised A17 OPMD mice. In the resistance overload protocol, we also observed an increased collagen deposition in the A17 OPMD mice which was associated with larger muscle mass and fiber cross sectional area and increased absolute maximal force as well as a reduction in PABPN1 aggregate number.

**Conclusions:** Running exercise and mechanical overload led to very different outcome in skeletal muscles of A17 mice. Both types of exercise enhanced collagen deposition but while the running protocol increased aggregates, the OVL reduced them. More importantly OVL reversed muscle atrophy and maximal force in the A17 mice. Our study performed in a relevant model gives an indication of the effect of different types of exercise on OPMD muscle which should be further evaluated in humans for future recommendations as a part of the lifestyle of individuals with OPMD.

## INTRODUCTION

Oculopharyngeal muscular dystrophy (OPMD) is a late-onset autosomal dominant inherited neuromuscular disorder, characterized by progressive eyelid drooping, swallowing difficulty and proximal limb weakness in the late stages of the disease. This disease is caused by mutations in the *PAPBN1* gene, consisting in a short (between 1 and 8) and meiotically stable GCN trinucleotide repeat expansion in its coding region^1^. The translation of the PABPN1 allele containing expanded repeats leads to a longer polyalanine tract in PABPN1 inducing - through protein misfolding - the formation of PABPN1 intranuclear aggregates^2^ in muscle fibers, the main hallmark of OPMD. In addition to PABPN1, these aggregates contain several proteins, such as ubiquitin, proteasome subunits, heat-shock proteins, splicing factors, poly(A) polymerase (PAP) as well as poly(A) RNA^3–5^. No treatment is currently available for the pathology. Preclinical studies in the A17 mouse model of OPMD using anti-aggregation drugs^6,7^ or gene therapy^8,9^ have shown beneficial effects.

Both endurance and resistance chronic exercises have been shown to have beneficial effects on skeletal muscle homeostasis both in aging and in pathological conditions. For example, treadmill running accelerated muscle repair and muscle stem cell function in old mice, through modulation of molecular pathways such as the upregulation of CyclinD1 expression and repression of Transforming Growth Factor-β (TGFβ) signaling^10^. Chronic exercise in the elderly was also shown to be beneficial to improve insulin sensitivity^11^, mitochondrial function^12^ and muscular regenerative capacity^13^. The favorable effects of physical activity on muscles have also been studied in several models of muscle disease and confirmed in different muscles. For example, the *mdx* mouse, a murine model of Duchenne Muscular Dystrophy (DMD), presented an increase in soleus muscle maximal force and a reduction in fatigue in both soleus and plantaris muscles following 16 weeks of voluntary wheel running^14^. Similarly, a 10-week swimming exercise increased the specific maximal force (absolute maximal force relative to muscle size) in the extensor digitorum longis (EDL) and soleus muscles of aged *mdx* mice^15^. The absolute and specific forces as well as fatigue resistance were also improved in tibialis anterior (TA) muscle of *mdx* mice by a 24-week treadmill running protocol with 3 exercises/week^16^. Moreover, long-term (1 year) voluntary running counteracted age-associated loss in absolute maximal force in *mdx* soleus muscle and reduced the fatigability in the EDL by 9%^17^. These benefits were also observed as early as one week following voluntary wheel running with a reduced muscle fragility in *mdx* TA muscles^18^. In the myotonic dystrophy 1 (DM1) mouse model, treadmill running improved motor performance, forelimb grip strength and endurance, releasing Muscleblind-like 1 (MBNL1) protein from myonuclear foci and improving mRNA alternative splicing^19^. Mechanical overload, which is used as a model of resistance exercise training, induced muscle hypertrophy and increased absolute maximal force production as well as fatigue resistance in healthy and *mdx* mice^20,21^. This overload protocol also successfully improved plantaris muscle mass of 20-month old healthy mice^22^.

Interestingly, OPMD muscles show molecular and histological signs of premature muscle aging^23^. A recent OPMD case report suggested the use of rehabilitation exercise for OPMD patients to improve swallowing ability after diagnosis of progressive dysphagia and associated choking events^24^. However, no study has ever looked at the effect of exercise on OPMD muscles. In order to address this, we have used the A17 mouse model, widely employed for preclinical studies^6,7,9^ as it recapitulates some of the key features of OPMD such as muscle atrophy, PABPN1 aggregates and muscle fibrosis^25,26^. We assessed the effect of exercise using two protocols: the first is a treadmill running-based protocol used as an endurance exercise^27^ while the second is the mechanical overload of the plantaris muscle by synergist ablation of the soleus and the medial and lateral parts of the gastrocnemius muscles, as a model of resistance exercise training^28^. We expected that such endurance or resistance exercises might be beneficial and could reverse some of the pathological features of the OPMD muscles such as muscle atrophy and weakness, fibrosis or PABPN1 nuclear aggregates. Some of these features are also observed in aging and in different muscle diseases. However, while the first endurance protocol produced no major effects on the OPMD phenotype, the second resistance protocol induced improvements with an increase in muscle mass, muscle fiber cross sectional area (CSA) and absolute maximal force as well as reduction in PABPN1 nuclear aggregates. Altogether this study gives us an indication of the effect of exercise on OPMD muscle which should be further evaluated in humans in order to provide future recommendations as a part of the lifestyle plan for patients with OPMD.

## MATERIAL AND METHODS

### Mice

A17 transgenic mice have been previously described^25,26^. Male A17 and FvB littermates were generated by crossing the heterozygous carrier strain A17 with FvB mice. The mice were genotyped by PCR^25^. Animals were housed with food and water ad libitum in minimal disease facilities (UMS28, Sorbonne Université, Paris). All experimentations in animals were performed in accordance with the French and European Community legislation (approval no. APAFIS #29361-20211012718186380 v3).

### Exercise protocols

Treadmill running protocol: A17 and FvB male mice followed an exercise training based on an established protocol (TREAT-NMD, SOP number: DMD_M.2.1.001). Groups of 5 FvB mice (12-13 weeks old) and 6 A17 mice (12-13 weeks old) were kept sedentary while groups of 5 FvB mice and 10 A17 mice were exposed to a running protocol. During 6 weeks, 3 days per week, FvB and A17 mice had a warm up time (0 to 20 cm/s during 2 min) followed by the exercise session (running on a treadmill for 20 min at 20 cm/s). TA and gastrocnemius muscles of both legs were collected 3 days after the end of the exercise protocol and compared to controls.

Mechanical overload protocol: To mimic resistance training, we performed the overload protocol by surgically removing the soleus muscle as well as the major distal parts of the lateral and medial gastrocnemius muscles in both legs of anesthetized mice (isoflurane 3%). This leads to mechanical overload of the plantaris muscles, as previously described^20,21,29^. Buprenorphine (Vetergesic®, 0.1 mg/kg) was administrated twice a day during 3 days after the surgery. Plantaris muscles of both legs were collected 1 month after the surgery and compared to control. 5 A17 and 6 FvB 42-week-old mice were subjected to overload protocol and compared to 6 A17 and 7 FvB mice that did not undergo surgery.

### *In situ* muscle function

Tibialis anterior (TA) and plantaris muscle functions were evaluated by measuring *in situ* isometric force, as previously described^30,31^. Briefly, mice were anesthetized and the distal tendon of the TA or plantaris muscles were cut and connected to an isometric transducer (Harvard Apparatus) using 4.0 braided surgical silk (Interfocus, Cambridge, UK). The foot was secured to a platform and the ankle and knee immobilized using stainless steel pins. Then, the sciatic nerve was exposed and stimulated by a bipolar silver electrode, using supramaximal square wave pulses of 0.1 ms duration. Labchart and Powerlab (hardware) from ADInstruments were used for data acquisition. We determined the optimal muscle length (Lo) by incrementally stretching the muscle using micromanipulators until the maximal isometric tetanic force was obtained. After 1 minute of muscle rest, the maximal isometric tetanic force (P0) was determined. We calculated the specific force (*g*/mg) by dividing the Po by TA or plantaris muscle weight. We let the muscle rest for 1 minute before the fatigue resistance measurement. For this, the muscle was continuously stimulated at 50 Hz for 20 seconds to obtain a submaximal continuous tetanus. The time necessary to obtain a force decrease of 30% was recorded. After the measurements, mice were euthanized by cervical dislocation.

### Samples harvesting, processing and storing

Animals were all euthanized at the end of the study or after the force measurement. TA and gastrocnemius (treadmill running protocol) or plantaris (overload protocol) muscles were harvested. Muscles were mounted on tragacanth gum (6% in water; Sigma-Aldrich) placed on a cork support and snapped frozen in liquid nitrogen-cooled isopentane for further RNA, histological and Western Blot analyses. All samples were stored at −80°C before analysis.

### Immunostaining and histological analyses

The muscle sections were prepared at 5 µm thickness, using a cryostat (Leica Biosystems, CM1850). and placed on coated glass slides (Thermofisher, Courtaboeuf, FR). Immunohistochemical staining was performed using the following antibodies: anti-PABPN1 (rabbit monoclonal, diluted 1:100, Abcam ab75855, overnight (ON), 4°C), anti-Dystrophin (Dys1) (mouse monoclonal, 1:20, clone Dy4/6D3 Novocastra, ON, 4°C). Alexa Fluor (Molecular Probes) antibodies conjugated to 488 or 555 nm were used. Briefly, after fixing sections in paraformaldehyde 4% during 1 hour (h) at room temperature (RT), the slides were incubated with KCl buffer (1M KCl, 30 mM HEPES, 65 mM PIPES, 10 mM EDTA, 2mM MgCl2, pH 6.9) for 1 h at RT. The muscle sections slides were blocked and permeabilized using 1% normal goat serum and 0,1% Triton X100 in PBS for 30 min at RT. Finally, the slides were incubated ON at 4°C with both PABPN1 and Dystrophin antibodies diluted in the blocking buffer. Slides were washed 3 times with PBS and incubated with fluorophore-conjugated secondary antibodies. The slides were washed 3 times with PBS, incubated 10 min at RT with Dapi (diluted 1:2,500 in PBS). Then they were washed 3 times with PBS and mounted using Dako mounting medium. 20x magnification images were randomly taken by a blinded observer and used to calculate the percentage of aggregates and CN fibers. A standard protocol for Sirius red and Hematoxylin Eosin were used to detect the collagen I and III and to observe the morphology of the muscle. NIH ImageJ analysis software was used to analyze the largest section of each muscle.

### Western blot

Proteins were extracted by sonication of the cryosection muscle tissue in RIPA buffer [0.1% deoxycholate sodium, 50 mM Tris-Hcl (pH8), 150 mM NaCl, 0.1 mM EDTA, 0.1% SDS and 1% NP40 with protease inhibitor cocktail (Complete, 11697498001, Roche Diagnostics) and phosphatase inhibitor cocktail (Santa Cruz; sc-45064)]. Protein concentration was determined by colorimetric detection method (Pierce BCA protein Assay, Thermo Fisher). Proteins were separated on 4-12% Bis-Tris gels (Invitrogen) and transferred onto a Polyvinylidene difluoride (PVDF) membrane for 1 h at constant 250 mA at 4°C. Membranes were blocked by incubation in 5% BSA in 1X TBS, 0.1% Tween-20 (TBS-T) for 1 h at RT under agitation. Following, membranes were stained with primary antibodies (PABPN1, Abcam, ab75855; GAPDH-HRP, Abcam) ON at 4°C under agitation. The next day, membranes were washed in TBS-T and incubated with appropriate secondary antibodies conjugated with HRP (except for GAPDH-HRP). The ChemiDoc Imaging System (Bio-Rad) was used to detect the signals from the membranes with Immobilon kit (Merck, WBKLS0500).

### RNA extraction

Total RNA was extracted from skeletal muscle samples using Trizol reagent (Invitrogen) according to the manufacturer’s instructions and Fast Prep Lysing Tube (MP Biomedicals). RNA concentration and quality were assessed with a NanoDrop® spectrophotometer ND-1000, and an Agilent 2100 bioanalyzer, respectively.

### RNA sequencing and analysis

After the paired end (2 ×75 bp) sequencing, a primary analysis based on AOZAN software (ENS, Paris) was applied to demultiplex and control the quality of the raw data (based ofFastQC modules / version 0.11.5). Obtained fastq files were then aligned using STAR algorithm (version 2.7.1a) on the GRCm38 reference from Ensembl, release 101, and quality control of the alignment realized with Picard tools (version 2.8.1). STAR parameters were the following: --sjdbOverhang 74 --twopassMode Basic -- outFilterType BySJout --quantMode TranscriptomeSAM. Reads were then counted using RSEM (v1.3.1) with the AlignedtoTranscriptome bam files and the statistical analyses on the read counts were performed with the R (v3.6.3) DESeq2 package version 1.26.0 to determine the proportion of differentially expressed genes between two conditions. During the statistical analysis, we filter out annotations where there are less than 3 samples with normalized counts greater than or equal to 10.

### Image acquisition and analysis

Images were visualized using a light (Leica DMR microscope equipped with a Nikon DS-Ri1 camera) or a fluorescent Olympus BX70 microscope (Olympus Optical, Hamburg, Germany) equipped with a CCD Camera (Photometrics CoolSNAP fx; Roper Scientific) and driven by Metaview (Universal Imaging, Downington, PA, USA)) microscope and analyzed using MetaMorph imaging system (Roper Scientific, Tucson, AZ, USA) software and ImageJ 1.440 for quantification analysis. CSA calculation was performed using MuscleJ plugin^32^.

### Graphical representation and statistical analysis

Results are presented as mean ± SEM. Statistical analysis performed using the GraphPad Prism 6.0 software. Unpaired Student t-test was used to compare two groups and one-way ANOVA with Tukey post-hoc test was performed to compare three groups or more. The threshold of significance was set at p<0.05.

## RESULTS

### Treadmill running induces no changes in OPMD muscle physiology

In order to determine if physical endurance training could affect the pathological phenotype of OPMD mice, both 12-13 weeks old FvB and A17 mice were subjected to an exercise protocol 3 times/week during 6 weeks (**Figure 1A**). At the end of the exercise protocol, no changes were observed in the body weight of the A17 and FvB mice (**Figure 1B**). Absolute maximal and specific maximal forces were analyzed in exercised mice and compared to those of FvB and A17 sedentary mice. As previously described^26^, A17 mice recapitulates the clinical OPMD phenotype compared to the FvB mice: reduced absolute maximal (p<0.0001) and specific maximal (p<0.0001) forces (**Figure 1C and 1D**) were associated with increased fatigue resistance (p=0.0302) of the TA muscle (**Figure 1E**). The physical endurance exercise protocol did not significantly modify absolute and specific maximal force or fatigue resistance in either FvB and A17 mice.

**Figure 1:**
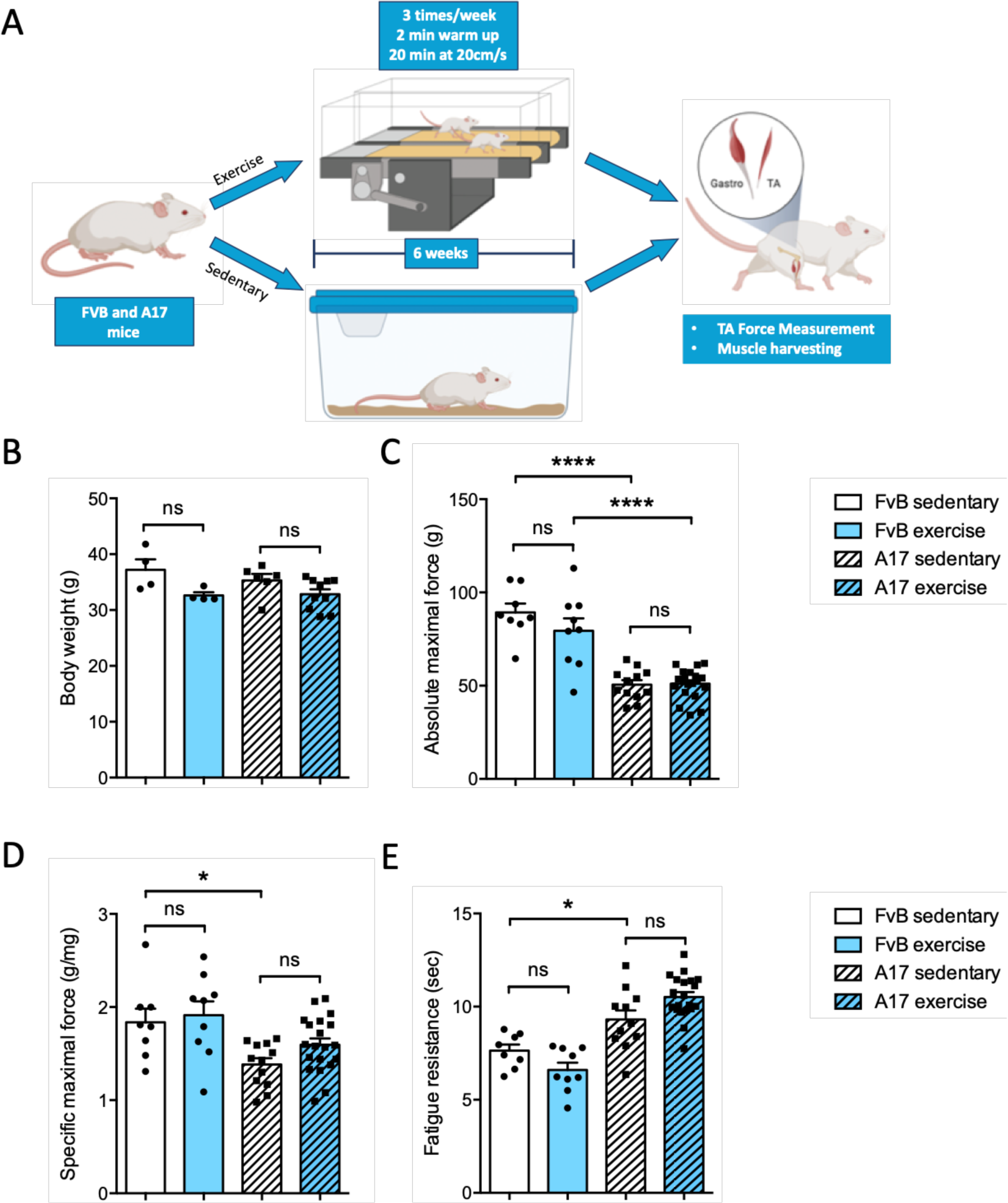
Treadmill running does not improve TA muscle physiology in OPMD mouse. (A) Schematic representation of treadmill protocol. (B) Final body mass (in grams, g) from sedentary or treadmill exercised FvB and A17 mice, n=4-10 mice/group. Absolute maximal force (D) (i.e. raw data without normalization) and specific force (E) (i.e. absolute maximal force (expressed in *g*) normalized to tibialis anterior (TA) muscle mass (in mg)) of TA muscle from sedentary or treadmill exercised FvB and A17 mice, n=8-20 muscles/group, (F) Fatigue resistance i.e. time to loss of 30% of the initial force (in sec) of TA muscle from sedentary or treadmill exercised FvB and A17 mice, n=8-20 muscles/group. ANOVA one-way followed by post-hoc Tukey multiple comparisons test, ns not significant, *p<0.05, ****p<0.0001. **Figure 1A** created with BioRender.com.

### Treadmill running protocol induces no change in OPMD muscle atrophy

To further assess the effects of our treadmill running protocol on muscle fibers, we analyzed the myofiber CSA of TA muscle. As shown on the representative pictures (**Figure 2A**), A17 TA muscles contain more small fibers compared to FvB control mice (**Figure 2B**) (p=0.0001), although the global fiber CSA mean was not significantly affected by the treadmill running protocol in A17 nor FvB mice (**Figure 2C**). Nevertheless, the TA weight was found decreased in A17 mice compared to FvB as previously described^26^ (p=0.0025) while treadmill running did not counteract this atrophy (**Figure 2D**).

**Figure 2:**
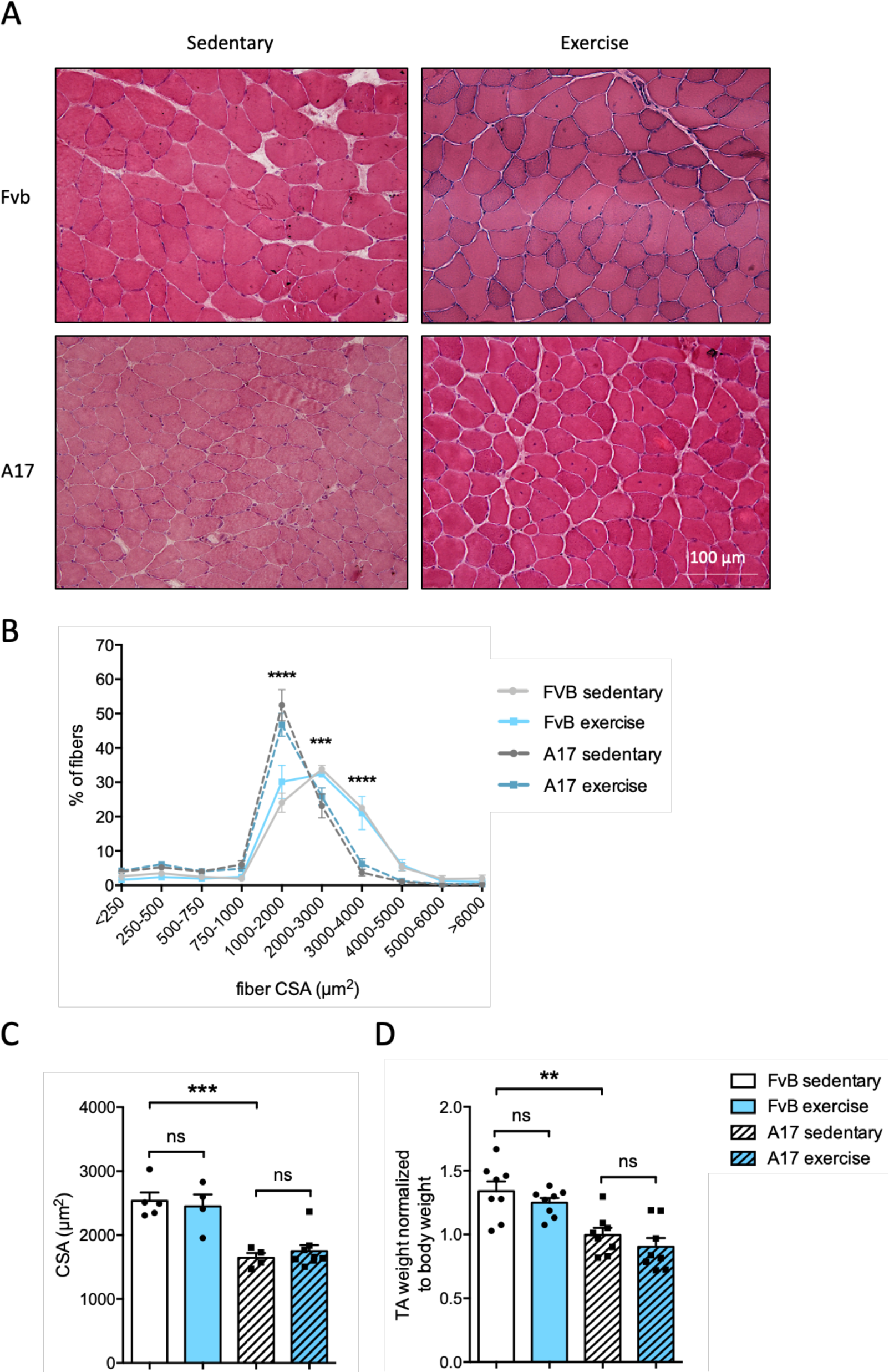
Treadmill running does not improve OPMD muscle atrophy. (A) Representative pictures of immunofluorescence staining of tibialis anterior (TA) muscle sections from sedentary or treadmill exercised FvB and A17 mice with dystrophin-1 staining (red) and nucleus (Hoechst, blue), magnification 20x. (B) Percentage of muscle fiber according to their cross-sectional area (CSA), from TA of sedentary or treadmill exercised FvB and A17 mice, n=4-10 mice/group, ANOVA two-ways followed by post-hoc Tukey multiple comparisons test, ***p<0.001, ****p<0.0001 between FvB sedentary and A17 sedentary groups. (C) Cross-sectional area (CSA) (in µm^2^) of TA muscles fibers from sedentary or treadmill exercised FvB and A17 mice, n=4-10 mice/group, (D) TA muscle mass (in mg) normalized to body mass (in g) from sedentary or treadmill exercised FvB and A17 mice, n=8 muscles/group. Panel C and D, ANOVA one-way followed by post-hoc Tukey multiple comparisons test, ns not significant, **p<0.01, ***p<0.001.

### PABPN1 nuclear aggregates are increased by treadmill running in OPMD mice

Accumulation of insoluble PAPBN1 aggregates in the nuclei of myofibers is a characteristic hallmark of OPMD^2^. In A17 mice, running increased the number of PAPBN1 intranuclear aggregates in the TA muscle (**Figure 3A-B**) by 24% (p=0.0066) without any change in PAPBN1 expression (**Figure 3C**). A similar increase in nuclear aggregates was also observed in gastrocnemius muscle **(Supplemental Figure 1.A**) (p<0.001). A RNAseq analysis performed on TA muscles of the four groups showed no change between sedentary and exercised groups as revealed by heatmap and PCA analysis (**Supplementary Figure 2**).

**Figure 3:**
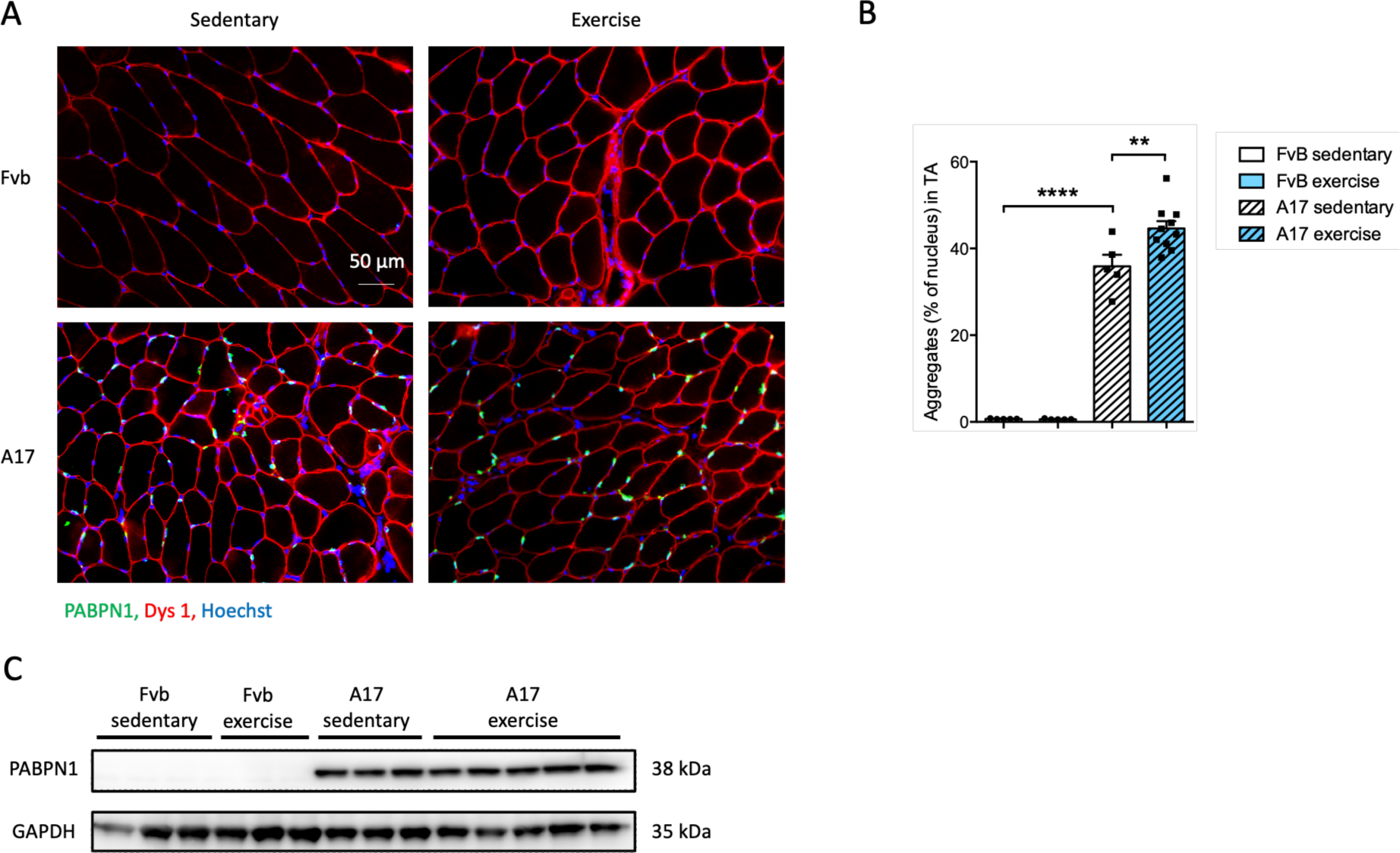
Treadmill running increases PABPN1 nuclear aggregates in OPMD TA muscle. (A) Representative pictures of immunofluorescence staining of tibialis anterior (TA) muscle sections from sedentary or treadmill exercised FvB and A17 mice with dystrophin1 (Dys1, red), PABPN1 (green) and nucleus (Hoechst, blue), magnification 20x. (B) Percentage of myonuclei containing a PABPN1 positive aggregate in TA muscle from sedentary or treadmill exercised FvB and A17 mice, n=3-9 mice/group, ANOVA one-way followed by post-hoc Tukey multiple comparisons test, **p<0.01, ****p<0.0001. (C) Representative western-blot of PABPN1 protein amount in TA muscles from sedentary or treadmill exercised FvB and A17 mice, GAPDH is used as loading control, n=3-5 mice/group.

**Supplementary Figure 1:**
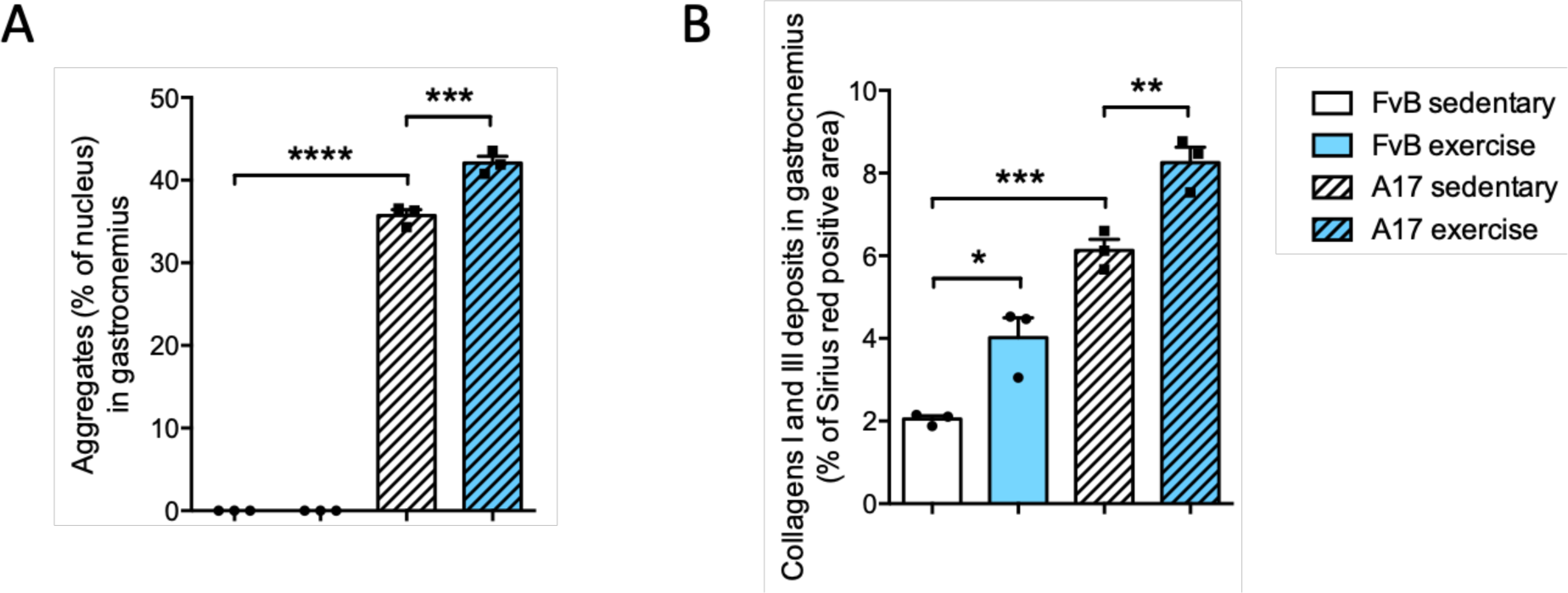
Treadmill running increases PABPN1 nuclear aggregates and collagen deposition in OPMD gastrocnemius muscle. (A) Percentage of myonuclei containing a PABPN1 positive aggregate in gastrocnemius muscle from sedentary or treadmill exercised FvB and A17 mice, n=3 mice/group. (B) Percentage of Sirius red positive area in gastrocnemius muscle from sedentary or treadmill exercised FvB and A17 mice. n=3 mice/group. ANOVA one-way followed by post-hoc Tukey multiple comparisons test, *p<0.05, **p<0.01, ***p<0.001, ****p<0.0001.

**Supplementary figure 2:**
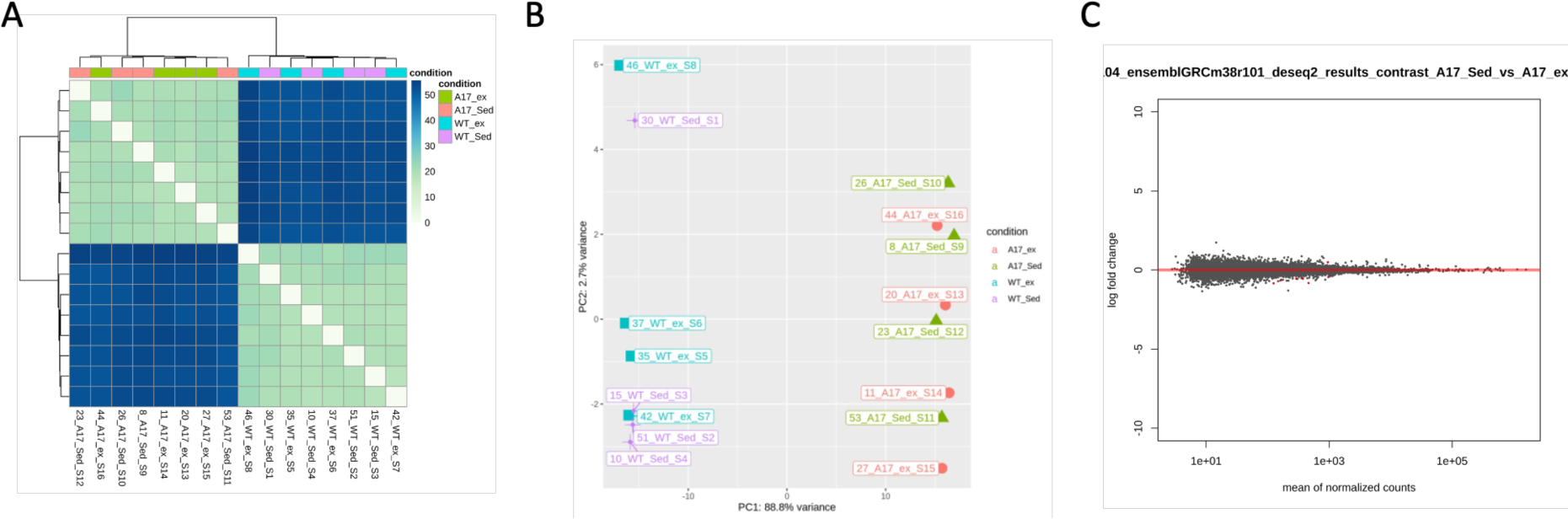
Treadmill training does not induce detectable gene expression changes in OPMD mouse TA muscle. (A) Heatmap of RNAseq tibialis anterior (TA) muscle samples, n=4 mice/group. (B) Principal component (PC) analysis heatmap appreciating the differences of global gene expression between FvB and A17 mice samples, n=4 mice/group. (C) Heatmap of differentially expressed genes between exercised A17 and sedentary A17 groups, black dot for non-significant expressed genes, red dot for significantly differentially expressed gene, n=4 mice/group.

### Treadmill running increases expression of collagen in OPMD mice

To assess the effects of a treadmill running on extracellular matrix markers, we performed a Sirius red staining, which detects mainly collagens I and III proteins (**Figure 4**). Sedentary A17 mice presented an increase in collagen level compared to sedentary FvB mice in the TA muscle (**Figure 4A-B**) (p=0.0387), which was further exacerbated by the treadmill protocol (p=0.0155). Collagen deposition was also enhanced in the gastrocnemius muscle in both FvB (p=0.0138) and A17 (p=0.009) mice compared to the respective sedentary mice (**Supplemental Figure 1.B**).

**Figure 4:**
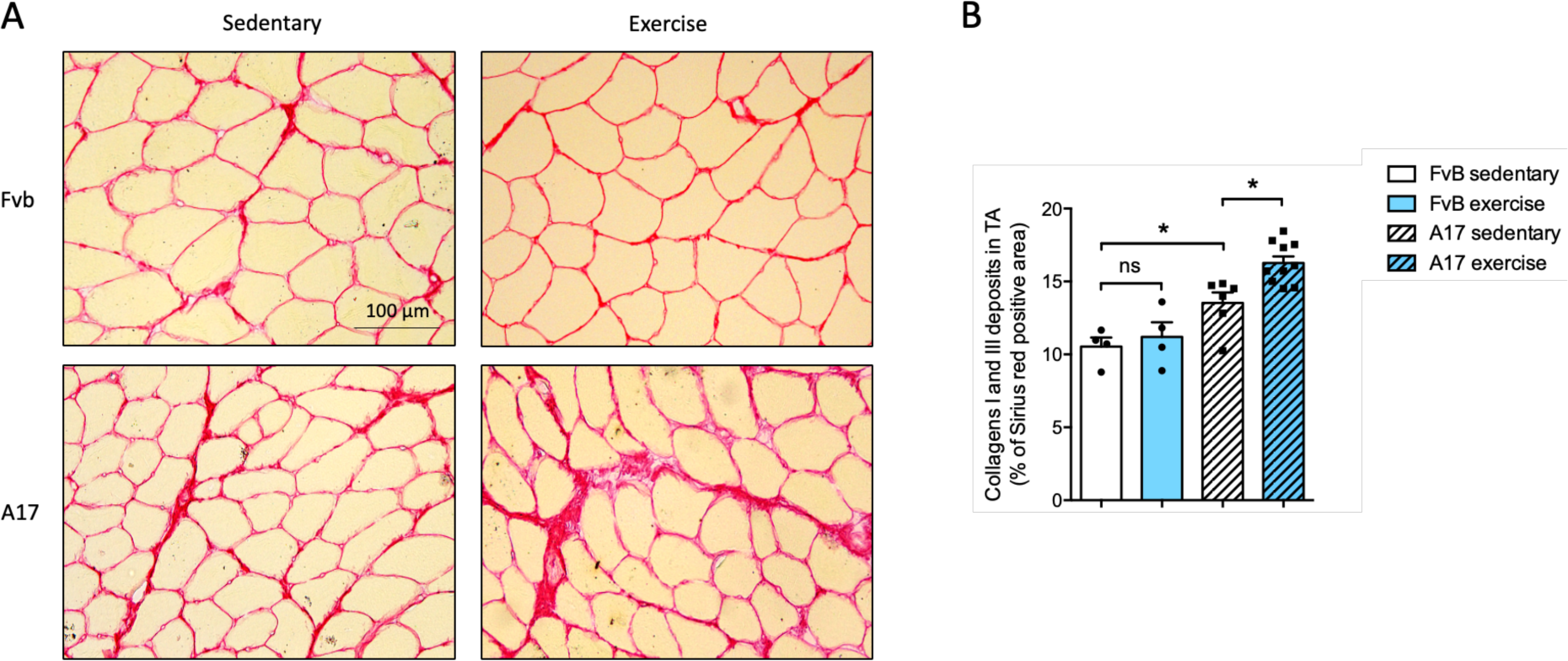
Treadmill training increases collagen deposition in OPMD TA muscle. (A) Representative pictures of Sirius red staining of tibialis anterior (TA) muscle sections from sedentary or treadmill exercised FvB and A17 mice, objective 20x. (B) Percentage of Sirius red positive area in TA muscle from sedentary or treadmill exercised FvB and A17 mice, n=3-9 mice/group, ANOVA one-way followed by post-hoc Tukey multiple comparisons test, ns: not significant, *p<0.05.

### Mechanical overload of plantaris muscle induces substantial changes in plantaris muscles of A17 mice

Since OPMD muscle phenotype shares similarities with muscle aging^23,33^, we decided to test a second protocol, mechanical overload (OVL) which mimics resistance training and has previously been shown to be beneficial to counteract muscle atrophy in aging muscle^22^ and muscle weakness in dystrophic murine muscle^18,20^. The model consists in ablating the major part of both the gastrocnemius and soleus muscles leading to mechanical overload of the plantaris muscle (**Figure 5A**)^34^. One month after surgery, plantaris muscles from treated A17 and FvB mice were collected and compared to the control groups.

**Figure 5:**
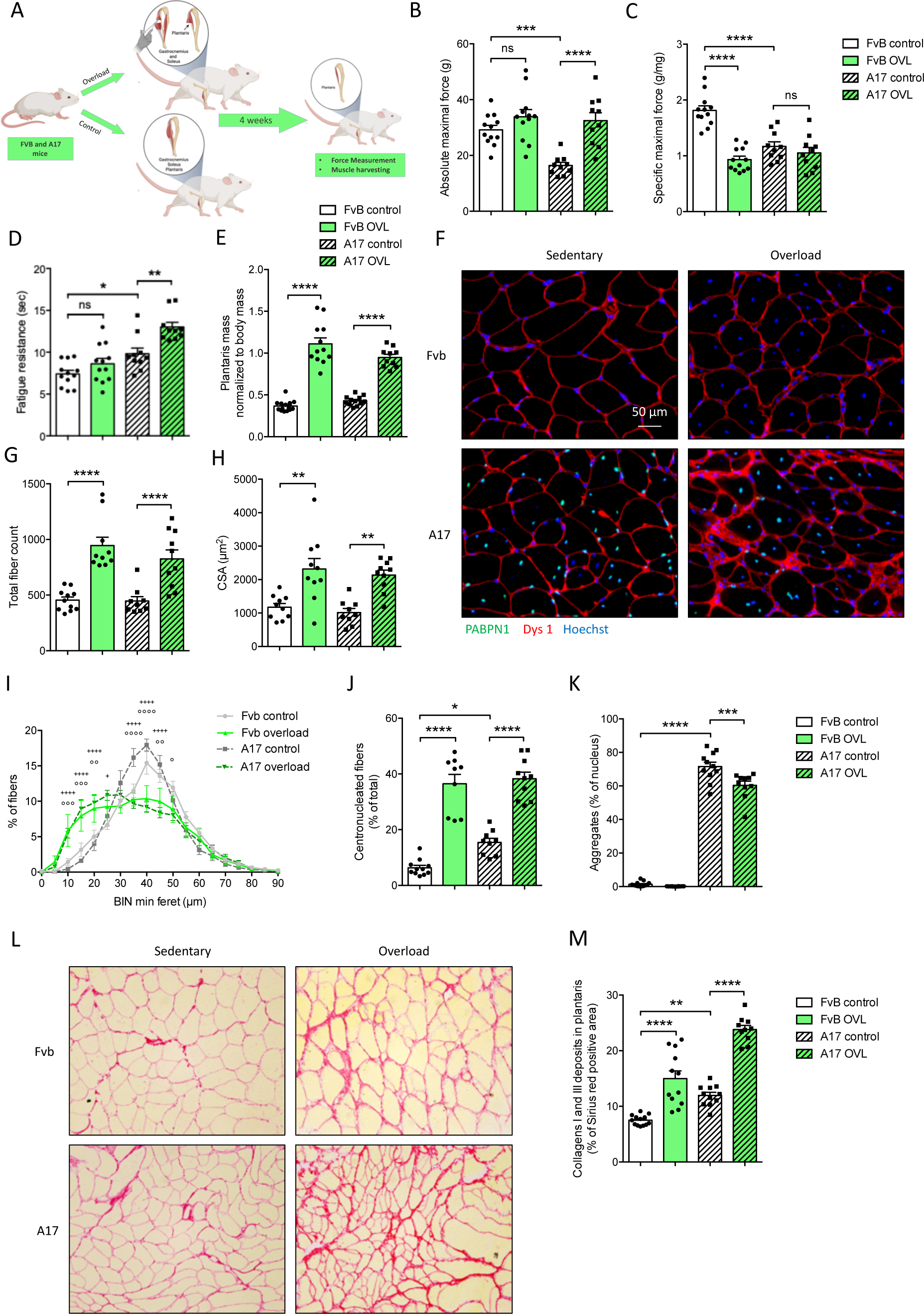
Mechanical overload shows beneficial effects in OPMD plantaris muscles. (A) Schematic representation of the overload protocol. Absolute maximal (B) and specific (C) maximal force of plantaris muscle from control or overload (OVL) FvB and A17 mice, n=10-12 muscles/group. (D) Fatigue resistance (time to lose 30% of the initial force, in sec) of plantaris muscle from control or OVL FvB and A17 mice, n=8-20 muscles/group, (E) Plantaris muscle mass (in mg) normalized to body mass (in g) from control or OVL FvB and A17 mice, n=10-14 muscles/group, (F) Representative pictures of immunofluorescence staining of plantaris muscle sections from control or overload FvB and A17 mice with dystrophin1 (red), PABPN1 (green) and nucleus (Hoechst, blue), objective 40x. (G) Total fiber count in plantaris muscle cross-section from control or OVL FvB and A17 mice, n=10-14 muscles/group, (H) Cross-sectional area (CSA) of plantaris muscles from control or OVL FvB and A17 mice, n=9-10 muscles/group, (I) Percentage of muscle fibers according to their cross-sectional area (CSA), from plantaris muscles of control or overload FvB and A17 mice, n=10-12 muscles/group, ANOVA two-ways followed by post-hoc Tukey multiple comparisons test, no differences between FvB control and A17 control groups, +p<0.05, ++++p<0.0001 between A17 control and A17 OVL groups, °p<0.05, °°p<0.01, °°°p<0.001, °°°°p<0.0001 between Fvb control and Fvb OVL groups. (J) Percentage of centro-nucleated fibers in plantaris muscles from control or OVL FvB and A17 mice, n=9-11 muscles/group, (K) Percentage of myonuclei containing a PABPN1 positive aggregate in plantaris muscle from control or OVL FvB and A17 mice, n=10-11 muscles/group, ANOVA one-way followed by post-hoc Tukey multiple comparisons test, (L) Representative pictures of Sirius red staining of plantaris muscle sections from control or overload FvB and A17 mice, objective 40x. (m) Percentage of Sirius red positive area in plantaris muscle from control or OVL FvB and A17 mice, n=11-12 muscles/group. For panels A-H, J and M, ANOVA one-way followed by post-hoc Tukey multiple comparisons test, *p<0.05, **p<0.01, ***p<0.001, ****p<0.0001. **Figure 5A** created with BioRender.com.

Sedentary plantaris muscles of A17 mice showed reduced absolute (**Figure 5B**) (p=0.0008) and specific (**Figure 5C**) (p=0.0252) maximal forces, as well as increased fatigue resistance (**Figure 5D**) (p=0.0287) and no changes in muscle weight (**Figure 5E**) compared to sedentary FvB mice. At the histological level, we observed an increase in the number of PABPN1 nuclear aggregates (**Figure 5F, 5K**) (p<0.0001) and an increased collagen deposition (**Figure 5L-M**) (p=0.0027) as expected^26^.

OVL increased absolute maximal force only in A17 mice (**Figure 5B**) (p<0.0001) and completely rescued this parameter to the sedentary FvB mouse level while specific maximal force was reduced by OVL in FvB mice only (p<0.0001) and unchanged in A17 (**Figure 5C**). Interestingly, fatigue resistance was improved in A17 mice specifically (**Figure 5D**) (p=0.0033). Muscle weight normalized to the body weight was increased by OVL (**Figure 5E**) in both the FvB (p<0.0001) and A17 mice (p<0.0001), as well as total fiber number per muscle cross section (**Figure 5G**) (p<0.0001 in FvB, p=0.0006 in A17). Consistently, fiber CSA was equally increased by the OVL protocol in FvB (p=0.0011) and A17 (p=0.0018) mice (**Figure 5F, 5H**). We also observed that OVL increased the percentage of centrally-nucleated fibers (CNF) in plantaris muscles of both FvB (p<0.0001) and A17 (p<0.0001) mice (**Figure 5I**) and reduced the percentage of myonuclei containing PABPN1 aggregates in A17 plantaris muscle (**Figure 5F, 5K**) (p=0.003). Finally, collagen deposition was increased in OVL plantaris muscles of exercised A17 (p<0.0001) and FvB (p<0.0001) mice (**Figure 5L-M**). Overall, OVL induced more significant changes in the muscles when compared to the treadmill running, suggesting an improved phenotype associated with reduction in aggregates and increased muscle mass and function.

## DISCUSSION

Numerous studies in rodent animal models as well as in humans have shown that physical exercise, either based on endurance or resistance, is beneficial in aging and in a number of muscular dystrophies. Firstly, in the context of aging, it has been shown that age-associated loss in force production and fatigability in soleus and EDL muscles in mice were improved by one year of voluntary running^17^. Resistance training achieved by mechanical overload protocol was also efficient as it increased the plantaris muscle mass in 20-month old mice^22^. Global beneficial effects of physical exercise have also been widely studied in older humans (for a review see^35^). Since OPMD muscles show many signs of premature muscle aging^23,33^, protocols applied to aging muscle which increase the muscle mass and strength could potentially be of interest for treating OPMD patients. Moreover, a recent study has revealed a loss of more than 80% in muscle strength, mobility and fatigue resistance in elderly OPMD patients^36^ justifying even more the clinical relevance of developing strategies to improve muscle function in OPMD patients.

Exercise protocols have also been applied in several muscle dystrophies, again both in mouse models and in patients. For example, voluntary exercise on a running wheel for 7 weeks reduced the number of RNA foci, while improving the muscle function and endurance in a mouse model of myotonic dystrophy type 1^19^. Interestingly, strength training during 12 weeks, using work out machines and aiming to increase limb muscle force, in myotonic dystrophy type 1 patients was found to counteract the alterations in skeletal muscle characteristics^37^. Positive effects have been widely described as consequence of chronic exercise in the mouse model of Duchenne Muscular dystrophy (DMD), the *mdx* mouse^14,15,18,18,20^. Accordingly, a 12-week protocol of isometric leg exercise in young DMD affected boys generated an improvement of leg muscle strength and function, without signs of damage^38^. Interestingly, a more acute exercise was generally shown to be detrimental in *mdx* mice (for a review see^39^) likely due to the lack of dystrophin in the sarcolemma of dystrophic myofibres that makes the muscles extremely susceptible of tissue damage. These examples prove that specific exercise protocols are efficient in improving aspects of muscular dystrophies and suggest that their effects should also be investigated in OPMD.

In this study, cohorts of A17 and FvB mice were first subjected to a endurance treadmill exercise protocol originally designed for *mdx* mice^27^. *mdx* mice, characterized by sarcolemmal fragility, were severely affected by this type of physical exercise because of the mechanical stress generated by the exercise. Here we observed that muscle mass and function, CSA and global gene expression of the A17 mice were not significantly modified by the treadmill protocol. After endurance exercise, we only observed a slight increase in collagen deposition and in the number of PABPN1 aggregates in the myonuclei of A17 mice. The increase in collagens I and III deposition was detected both in FvB and A17 mice suggesting this to be a temporary, non-pathological, enlargement of the extracellular matrix (ECM) generated by muscle remodeling as already showed for other exercise protocols in normal individuals (^40,41^, for a review see^42^). We conclude that in contrast to what was observed in *mdx* mice^39^, this exercise protocol has no major adverse effects in the A17 mice. The difference between *mdx* and A17 mice is likely on the lack of *a priori* loss of muscle fiber integrity in OPMD compared to DMD muscles that are damaged by such a relatively low intensity protocol^25,26^.

Intranuclear PABPN1 aggregates are the main histological hallmarks of OPMD since their discovery in 1980 by Fernando Tomé and Michel Fardeau^2^. They are the consequence of misfolding of expanded PABPN1 protein and are known to sequester RNA and proteins^3–5^. We have previously shown that therapeutic approaches reducing fibrosis, atrophy and improving muscle force contraction, and so ameliorating the pathology, are strictly correlated with a reduction in the number of nuclear aggregates^7,8^. This supported the general consensus that approaches increasing aggregates would be detrimental for muscle cells. In the present study the treadmill exercise induced an increase in the number of PABPN1 aggregates suggesting that exercising OPMD muscles may have had an effect on cellular stress which could increase the formation of aggregates. However, this increase in PABPN1 aggregates during the running protocol did not lead to any detrimental effect on muscle function since there was no modification in the functional parameters that were measured. This suggested that either this increase in aggregates was not sufficient to further modify the muscle pathology (as revealed also by the unchanged RNAseq) or that such aggregates caused by a relatively short-term exercise have different properties compared with those physiologically generated by the pathology.

To complement the treadmill exercise, we performed a resistance training protocol using the OVL model. This OVL protocol produced beneficial effects in the A17 mouse muscles as shown by a decrease of about 16% of nuclear aggregates in A17 mice, as well as an improvement in absolute maximal force, muscle mass and fiber cross-sectional area (CSA). It should be noted that while the absolute maximal force was increased, the specific maximal force did not change as the muscle was also larger. This finding was similar to the outcome of a previous study where myostatin downregulation, obtained by systemic delivery of a monoclonal antibody, significantly increased the muscle mass and the absolute maximal force without affecting the specific maximal force^43^. Moreover, the previously described increase in myonuclei number by mechanical overload protocol^44^ could contribute to the reduction observed in nuclear aggregation.

Like in the treadmill protocol, we observed an increase in ECM deposition in the OVL plantaris muscle. Our data are consistent with previous studies reporting an increase in the amount of the non-contractile tissue, hydroxyproline muscle content and *procollagen 1* mRNA expression following the overload protocol^41^. Moreover, it is known that ECM-related gene expression is induced following both acute resistance^45^ and endurance exercise^46,47^. For example, collagens IA2, IIIA1 and IVA1 are in the top five most upregulated genes in the transcriptome of skeletal muscle in response to endurance and resistance exercise^48^. Interestingly, a concomitant increase in both ECM deposition and muscle force was observed by the overload protocol^40^ suggesting the non-negative effect of the ECM deposition in the exercise context.

OVL is also characterized by an important increase in the number of centronucleated fibers (CNF)^21^. This effect is supposed to be due to the associated muscle regeneration but also to the generation of split fiber as previously shown by applying the OVL protocol in a muscle satellite cell null mouse model^22^. In our experiment, following overload, the plantaris muscles of A17 mice presented 38% of CNF along with 16% reduction in the number of nuclei containing PABPN1 aggregates. Accordingly, we have previously shown that muscle regeneration was associated with a decrease on the percentage of nuclei containing PABPN1 aggregates in human biopsies^5^.

Overall, this is the first time that the effect of exercise on OPMD pathology has been studied. Physical exercise has been investigated as a treatment in humans against muscular dystrophies^39,49^ and found to be generally positive. Here we have shown that a low-intensity exercise did not induce major modifications in OPMD mice muscles. The high-intensity exercise induced an increase in muscle mass and functionality suggesting that this type of physical exercise might show some benefits in individuals with OPMD.

Our study performed in the A17 mouse model gives indication as to the effect of different types of exercise on OPMD muscle which should be further evaluated in humans for future recommendations as a part of the lifestyle of individuals with OPMD.

## Acknowledgments

This work was financed by INSERM, Sorbonne University, the Fondation Recherche Médicale (EQUIPE FRM EQU201903007784) and the Agence Nationale pour la recherche ANR N° ANR-18-HDHL-0002-05. Dr Alexis Boulinguiez was supported by internal pump-prime funding from Royal Holloway, University of London. We thank Bruno Cadot from the MyoIMAGE facility for imaging support. We thank Maria Kondili for the help with bioinformatic post-analysis. We thank Franck Letourneur and the GENOM’IC Core Facility at Institut Cochin for the RNAseq analysis. The authors of this manuscript certify that they comply with the ethical guidelines for authorship and publishing in the Journal of Cachexia, Sarcopenia and Muscle.

## Author Contributions

CT, AM and EN designed the study. AB, JD, LM, CR, JF generated the data. AB, JD, BC, LG HRM performed data analysis. ML and BG performed the endurance exercise training and force measurements. LA and BSP performed the RNAseq analysis. AF performed the OVL protocol. AB, BC, LG, AM and CT wrote a first draft of the manuscript. All authors were then involved in the writing and revising of the manuscript.

